# Cryopreservation of human pluripotent stem cell-derived cardiomyocytes is not detrimental to their molecular and functional properties

**DOI:** 10.1101/700849

**Authors:** Lettine van den Brink, Karina O. Brandão, Catarina Grandela, Mervyn P.H. Mol, Christine L. Mummery, Arie O. Verkerk, Richard P. Davis

## Abstract

Human induced pluripotent stem cell-derived cardiomyocytes (hiPSC-CMs) have emerged as a powerful platform for *in vitro* modelling of cardiac diseases, safety pharmacology, and drug screening. All these applications require large quantities of well-characterised and standardised batches of hiPSC-CMs. Cryopreservation of hiPSC-CMs without affecting their biochemical or biophysical phenotype is essential for facilitating this, but ideally requires the cells being unchanged by the freeze-thaw procedure. We therefore compared the *in vitro* functional and molecular characteristics of fresh and cryopreserved hiPSC-CMs generated from two independent hiPSC lines. While the frozen hiPSC-CMs exhibited poorer replating than their freshly-derived counterparts, there was no difference in the proportion of cardiomyocytes retrieved from the mixed population when this was factored in. Interestingly, cryopreserved hiPSC-CMs from one line exhibited longer action potential durations. These results provide evidence that cryopreservation does not compromise the *in vitro* molecular, physiological and mechanical properties of hiPSC-CMs, though can lead to an enrichment in ventricular myocytes. It also validates this procedure for storing hiPSC-CMs, thereby allowing the same batch of hiPSC-CMs to be used for multiple applications and evaluations.

## 1. Introduction

Human pluripotent stem cell-derived cardiomyocytes (hPSC-CMs) are now an established *in vitro* model and tool for studying cardiovascular development and disease, safety pharmacology and drug development, as well as having potential therapeutic applications (Brandão et al., 2017; Gerbin and Murry, 2015; Hartman et al., 2016; Magdy et al., 2018). Due to the significant progress made in efficiently differentiating hPSCs to cardiomyocytes, it is now possible to generate the large quantities of hPSC-CMs required for these purposes which can then be evaluated using a multitude of assays (Denning et al., 2016). To facilitate this, cryopreservation of hPSC-CMs is essential. Not only does the ability to freeze hPSC-CMs make the generation of these cells more cost and time effective, it also enables the same batch of hPSC-CMs to be analysed at multiple time points; thereby reducing this source of variability when performing multiple assays to investigate a disease phenotype or when performing large-scale drug discovery screens. From a clinical perspective, cryopreservation of the hPSC-CMs is also a necessary step to provide sufficient time for performing quality control checks (Fujita and Zimmermann, 2018). Furthermore, cryopreservation allows the hPSC-CMs to be easily distributed among users, including to laboratories without the expertise or infrastructure to culture and differentiate hPSCs. Indeed, there are now several commercial suppliers that distribute cryopreserved hPSC-CMs for both academic and commercial applications (Blinova et al., 2018; Kitaguchi et al., 2016; Maddah et al., 2015).

Several reports have described procedures for freezing hPSC-CMs and using the resulting cryopreserved hPSC-CMs, for example in transplantation studies or for the generation of engineered heart tissue (Breckwoldt et al., 2017; Chong et al., 2014). While these protocols vary in the composition of the freezing medium, the majority include 10% dimethylsulfoxide (DMSO) as a cryoprotective agent and require the hPSC-CMs to be enzymatically dissociated into single cells. Obviously, it is critical that the cryopreservation procedure does not alter the molecular, biochemical or functional phenotype of the hPSC-CMs. While it has been demonstrated that frozen hPSC-CMs are viable upon thawing, express cardiac-specific markers, exhibit typical electrophysiological, calcium handling and contractility characteristics, and can electrically couple in grafts (Blinova et al., 2018; Gerbin et al., 2015; Hwang et al., 2015; Kim et al., 2011; Puppala et al., 2013), these studies did not compare freshly-derived hPSC-CMs head-to-head with frozen-thawed batches. The few studies that have undertaken such a comparison have focussed on cardiomyocyte purity and viability immediately after thawing, as well as engraftment efficiency in rodents and non-human primates (Chen et al., 2015; Chong et al., 2014; Xu et al., 2011).

To date there have been no reports comparing the *in vitro* phenotype and physiological properties of cryopreserved and non-frozen hPSC-CMs following replating. Such studies are warranted due to the increasing use of hPSC-CMs for investigating disease mechanisms and in safety pharmacology assays (Brandão et al., 2017; Denning et al., 2016; Giacomelli et al., 2017). For this reason, we have evaluated freshly-derived and cryopreserved cardiomyocytes generated from two different human induced pluripotent stem cell lines (hiPSC-CMs). The hiPSC-CMs were compared in terms of replating and cardiomyocyte subtype, as well as their electrophysiological and mechanical properties. Although the cryopreserved cells exhibited poorer replating efficiency after thawing, when this was adjusted for, there was no difference in the proportion of cardiomyocytes recovered compared to the non-frozen cells. Similarly, there was no significant difference in the percentage of ventricular hiPSC-CMs. However, hiPSC-CMs from one line did show prolonged action potential (AP) duration in the cryopreserved compared to the fresh cardiomyocytes. Besides this, no other differences in the electrical and mechanical properties were observed, indicating that cryopreservation does not appear to be detrimental for the physiological attributes of hiPSC-CMs *in vitro*. As such, this study confirms that cryopreserved hPSC-CMs retain their *in vitro* molecular and functional characteristics and validates this as an opportune method for stockpiling hPSC-CMs for their use in downstream applications.

## 2. Results

### 2.1 Differentiation of hiPSCs to cardiomyocytes

Two independent hiPSC lines (LUMC20 and LUMC99) were differentiated into cardiomyocytes, with spontaneously contracting regions typically observed around day 8 (d8) of differentiation (**Fig. 1A and 1B**). Both cell lines efficiently generated cardiomyocytes, with on average 82.7%±7.6% (LUMC20) and 86.5%±3.0% (LUMC99) of cells expressing the pan-cardiomyocyte marker cardiac troponin T (cTnT) at d21 of differentiation (**Fig. 1C and 1D**). At d21, the hiPSC-CMs were dissociated and either immediately replated (*fresh*) or cryopreserved for at least 7 days (on average 28 days). Cryopreservation was performed by a rate-controlled (−1°C/minute) temperature decrease to −80°C in a freezing medium comprising 90% KnockOut Serum Replacement (KSR) and 10% DMSO. Non-frozen and thawed hiPSC-CMs from the same differentiation were compared under identical experimental conditions in terms of replating efficiency, cardiac marker expression, and biophysical characteristics (AP and contraction).

**Figure 1.**
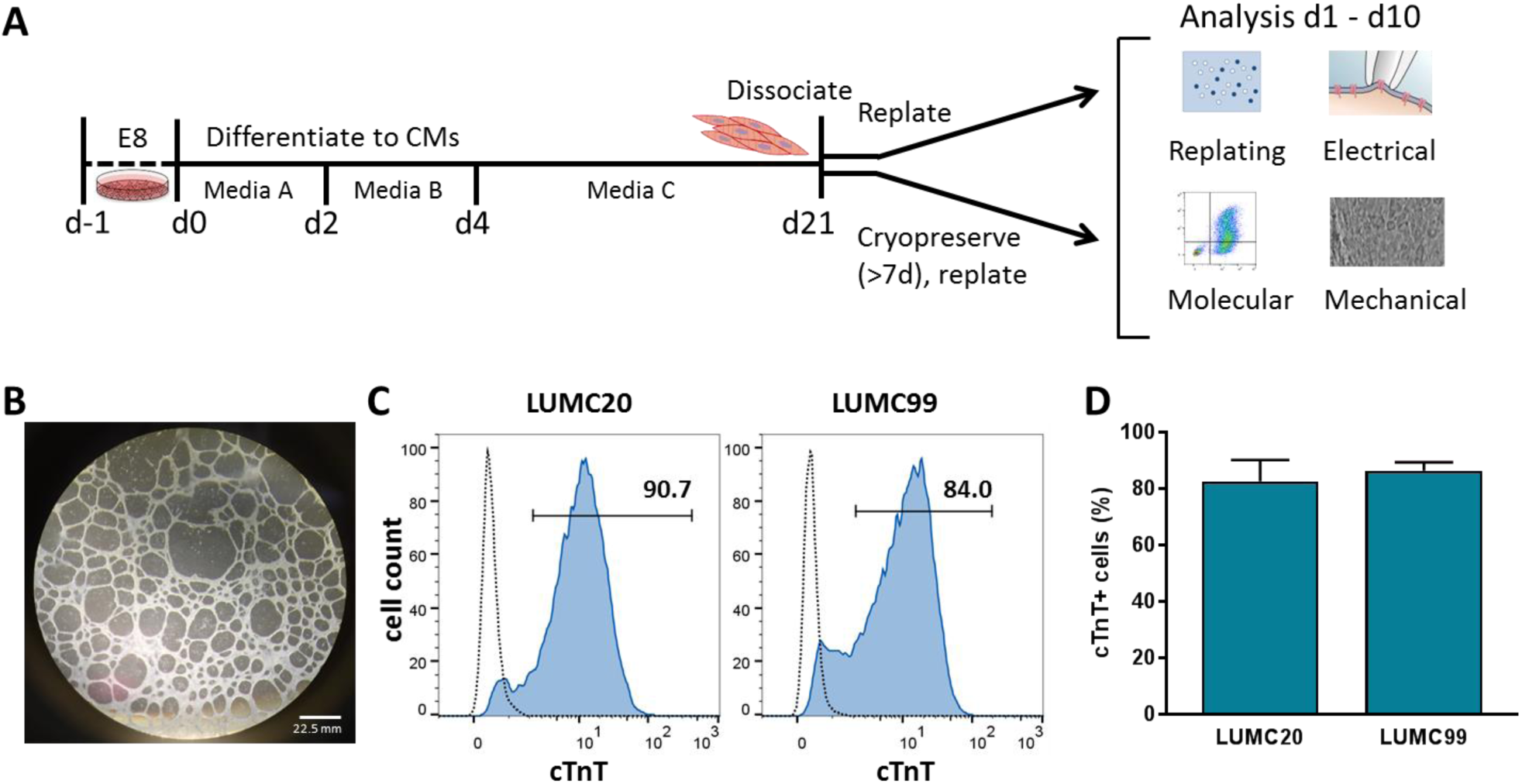
Differentiation of hiPSC lines LUMC20 and LUMC99 to cardiomyocytes prior to cryopreservation and replating. **A)** Schematic outlining the differentiation procedure and subsequent analyses performed. At differentiation day (d)21, the dissociated hiPSC-CMs were either directly replated, or cryopreserved for at least 7 days before being thawed and replated. **B)** Phase-contrast image of a well containing d20 hiPSC-CMs derived from LUMC20. **C)** Representative histogram plots showing the percentage of cTnT^+^ cells at d21 as determined by flow cytometry. Dotted lines represent a control cTnT^-^ population. **D)** On average, for both LUMC20 and LUMC99, more than 80% of the cultures were cardiomyocytes as determined by flow cytometry analysis of cTnT (n=7 and 4 respectively).

### 2.2 Cryopreservation affects cell survival of replated cultures but not the relative hiPSC-CM contribution

To evaluate whether cryopreservation had any effect on cell viability, dissociated hiPSC-CMs were stained with Trypan blue before and after freezing (**Fig. 2A**). There was no significant difference in the percentage of viable cells for LUMC20 (98.3±0.3% non-frozen vs 95.7±1.4% frozen, p=0.08; unpaired t-test), indicating that the cryopreservation procedure did not cause significant necrosis. While cryopreserved hiPSC-CMs attached and expressed the cardiac markers α-actinin and myosin heavy chain (**Fig. 2B**), 24h after replating the recovery of the cryopreserved cultures was approximately half that of their freshly replated counterparts (15.5×10^−3^ ± 1.9×10^−3^ A.U. non-frozen vs 9.2×10^−3^ ± 1.0×10^−3^ A.U. frozen per 1.0×10^4^ cells seeded) (**Fig. 2C**). This difference persisted for at least 7 days post-replating, suggesting that there was also no difference in proliferation rates between the differentiated cultures that were freshly replated or cryopreserved. Similar differences in replating efficiency were also observed for differentiated cells from LUMC99 (**Supplementary Fig. S1**).

**Figure 2.**
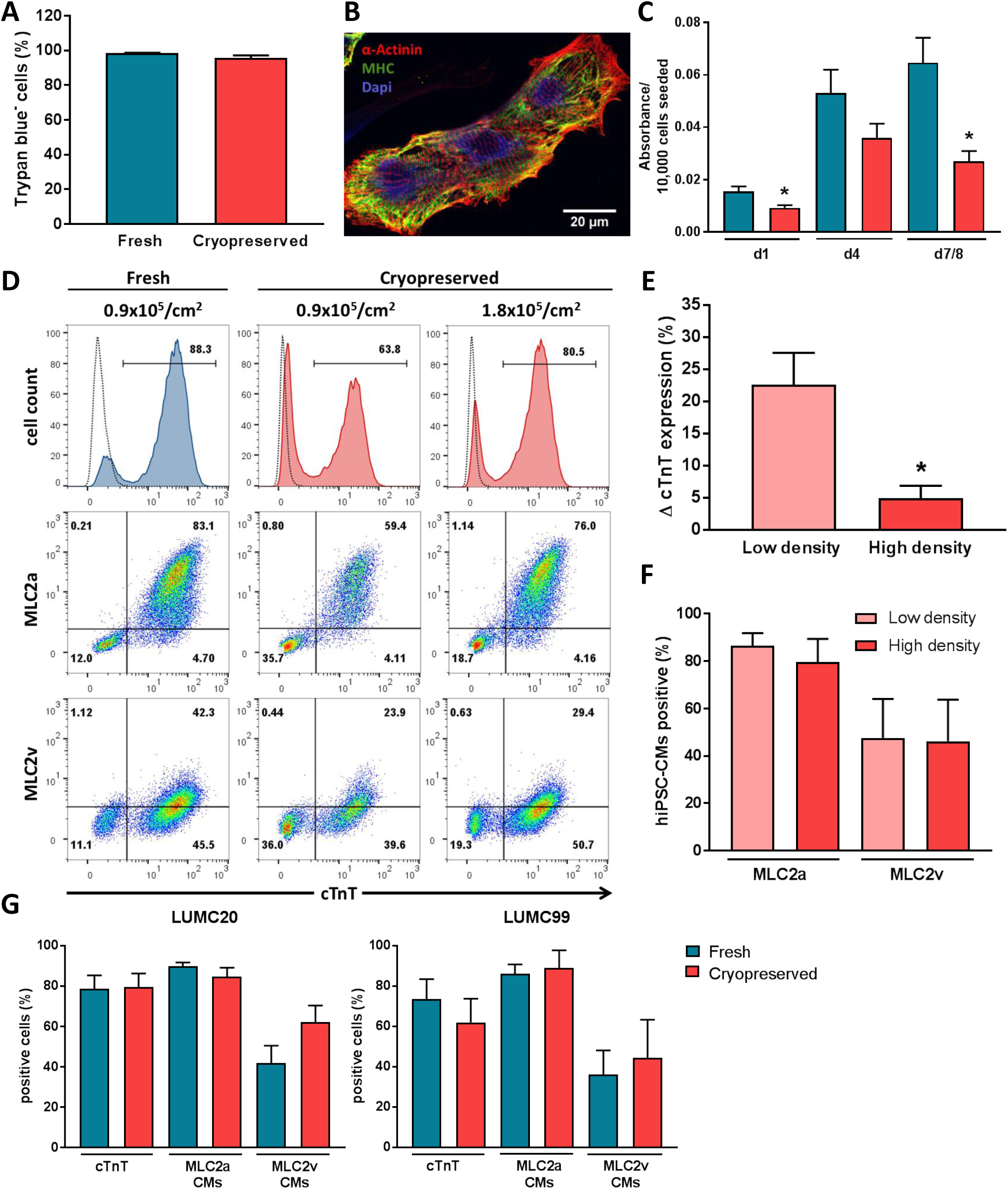
Replating recovery and cardiac marker expression of non-frozen and cryopreserved hiPSC-CMs derived from LUMC20. **A)** Percentage of viable cells directly after dissociation (*fresh*) or upon thawing (*cryopreserved*); n=14 from 7 independent differentiations. **B)** Immunofluorescence image of cryopreserved hiPSC-CMs 6 days after thawing following staining with the sarcomere markers, α-actinin (red) and myosin heavy chain (MHC, green). Nuclei were stained with DAPI (blue). **C)** Recovery of fresh (blue) and cryopreserved (red) hiPSC-CMs at day 1, 4 and 7/8 post-replating. * indicates statistical significance (day 1 p=0.011, day 7/8 p=0.007, unpaired *t*-test); n=8 from 4 independent differentiations **D)** Representative flow cytometric analysis of d21+7 hiPSC-CMs for expression of cTnT, MLC2a and MLC2v. Left column is non-frozen CMs, while the remaining columns are cryopreserved hiPSC-CMs plated at either the same density (0.9×10^5^/cm^2^; *centre column*) or two-fold higher density (1.8×10^5^/cm^2^; *right column*) as the non-frozen CMs. Top row depicts histogram plots of cTnT expression, while remaining rows depict bivariate density plots of MLC2a/cTnT (*middle row*) and MLC2v/cTnT (*bottom row*). Numbers inside the plots are percentage of cells within the gated region. Dotted lines represent a control cTnT^-^ population. **E)** Bar graph showing difference (D) in cTnT expression compared to the corresponding fresh hiPSC-CMs when the cryopreserved hiPSC-CMs were seeded at low (0.9×10^5^/cm^2^) or high (1.8-2.1×10^5^/cm^2^) density. * indicates statistical significance (p=0.016, unpaired *t*-test) (n=4) **F)** Bar graph showing percentage of cryopreserved cTnT^+^ hiPSC-CMs expressing MLC2a or MLC2v when seeded at either low (0.9×10^5^/cm^2^) or high (1.8×10^5^/cm^2^) density (n=4) **G)** Bar graph showing percentage of freshly replated and cryopreserved cells expressing cTnT, as well as proportion of cTnT^+^ hiPSC-CMs expressing MLC2a or MLC2v after taking into account differences in replating efficiencies for both LUMC20 (*left*) and LUMC99 (*right*) (n=7 and n=4 respectively).

We therefore assessed whether this difference affected the proportion of hiPSC-CMs present in the replated cultures by flow cytometric analysis for cTnT, as well as the atrial and ventricular myosin light chain (MLC) isoforms (MLC2a and MLC2v respectively). The replating density did not affect the proportion of hiPSC-CMs that recovered when the cells were freshly replated (**Supplementary Fig. S2A**), therefore the non-frozen hiPSC-CMs were subsequently always seeded at 0.9×10^5^/cm^2^. However for the cryopreserved cells, higher seeding densities improved hiPSC-CM recovery based on the proportion of cells expressing cTnT (**Supplementary Fig. S2B**), resulting in similar percentages to that observed in the corresponding non frozen hiPSC-CMs (**Fig. 2D and 2E**). Altering the seeding density also resulted in a corresponding increase in MLC2a^+^ and MLC2v^+^ cells (**Fig. 2D and Supplementary Fig. S2B**), although the overall percentage of hiPSC-CMs (cTnT^+^ cells) expressing these markers did not vary between the different seeding densities (**Fig. 2F**). This indicated that while higher seeding densities improved the recovery of cryopreserved hiPSC-CMs, it did not influence the cardiomyocyte subtype or maturity.

We therefore factored in the difference in replating efficiency between the non-frozen and cryopreserved hiPSC-CMs by seeding approximately twice as many cryopreserved hiPSC-CMs per cm^2^. We then observed no difference in the percentage of cTnT^+^ hiPSC-CMs in the replated populations for the LUMC20 line (78.6±6.8% non-frozen vs 79.6±6.7% frozen), as well as for the LUMC99 line (73.4±10.0% non-frozen vs 61.9±11.8% frozen) (**Fig. 2G**). Similarly, there was no difference in the proportion of hiPSC-CMs expressing MLC2a (LUMC20, 89.8±1.9% vs 84.7±4.4%; LUMC99, 86.1±4.6% vs 89.0±8.7%; non-frozen vs frozen) (**Fig. 2G**). Although not significant, we did observe a trend towards a higher percentage of hiPSC-CMs expressing MLC2v following cryopreservation for both cell lines (LUMC20, 41.7±8.8% vs 62.1±8.2%; LUMC99, 36.1±12.0% vs 44.3±19.0%; non-frozen vs frozen) (**Fig. 2G**), suggesting that cryopreservation might promote the maturation of hiPSC-CMs to a ventricular subtype. Prolonged storage of the hiPSC-CMs did not affect expression of these cardiac markers, with the same batch of cells having similar values at both 1- and 8-months post-cryopreservation (**Supplementary Fig. S3**). Taken together, these results indicated that while frozen hiPSC-CMs exhibited poorer replating efficiencies, cryopreservation did not significantly affect the proportion of hiPSC-CMs or subtypes recovered after both short-and long-term storage. However, an increase in the proportion of hiPSC-CMs expressing MLC2v was observed compared to the non-frozen counterparts.

### 2.3 Cryopreservation is not detrimental to the electrical activity of hiPSC-CMs

Next, we performed AP measurements of fresh and cryopreserved hiPSC-CMs generated from the same differentiation. APs were recorded from single spontaneously contracting hiPSC-CMs, and to obtain a close-to-physiological resting membrane potential (RMP) we injected an in silico inward rectifier K^+^ current (I_K1_) with Kir2.1 characteristics using dynamic clamp methodology (Meijer van Putten et al., 2015). **Fig. 3A** shows representative APs from frozen and non-frozen cardiomyocytes derived from the two hiPSC lines, and average AP characteristics are summarized in **Fig. 3B** (LUMC20) and **Fig. 3C** (LUMC99). Neither maximum AP upstroke velocity (V_max_) nor AP amplitude (APA) showed significant differences between fresh and cryopreserved hiPSC-CMs from both hiPSC lines. The RMP as well as AP duration (APD) at 20, 50, and 90% repolarisation (APD_20_, APD_50_, and APD_90_, respectively) were unaffected by cryopreservation for hiPSC-CMs derived from the LUMC99 hiPSC line (**Fig. 3C**). However, cryopreserved hiPSC-CMs from the LUMC20 hiPSC line showed a hyperpolarised RMP and prolonged APD (**Fig. 3B**). While differences in RMP might affect APDs by altering availability of sodium, calcium and potassium currents (Verkerk et al., 2017), we found no correlation between APD_90_ and RMP (data not shown), suggesting that the longer APs in the frozen LUMC20 hiPSC-CMs may be due to the greater proportion of MLC2v^+^ cardiomyocytes. Nevertheless, these results indicated that cryopreservation is not detrimental to the electrophysiological properties of the hiPSC-CMs *in vitro*.

**Figure 3.**
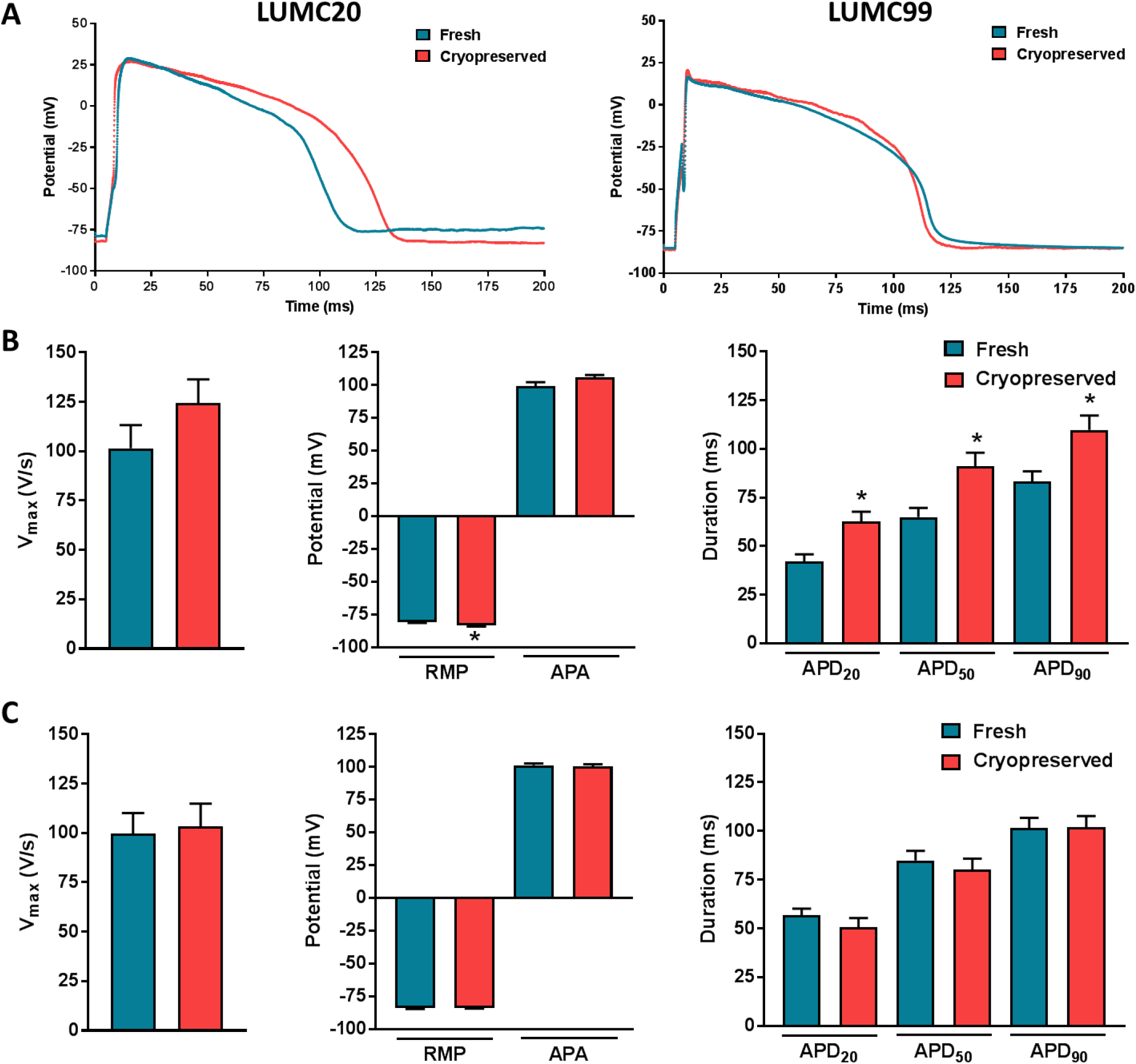
Action potential (AP) characteristics of non-frozen and cryopreserved hiPSC-CMs. **A)** Representative AP traces of fresh and cryopreserved hiPSC-CMs measured at 1 Hz for both LUMC20 and LUMC99 cell lines. **B & C)** Average data at 1 Hz for maximal upstroke velocity (Vmax), resting membrane potential (RMP), AP amplitude (APA) and AP duration at 20, 50, and 90% of repolarization (APD20, APD50, and APD90, respectively) for fresh and cryopreserved hiPSC-CMs for both LUMC20 (**B**) and LUMC99 (**C**) cell lines. For **B**, n=34 and n=38 respectively from 4 independent differentiations; for **C**, n=45 and n=40 respectively from 3 independent differentiations. * indicates statistical significance (RMP p<0.001, APD20 p=0.002, APD50 p=0.003, APD90 p=0.006 unpaired t-test)

### 2.4 Frozen and non-frozen hiPSC-CMs display similar contraction characteristics

Finally, we investigated whether cryopreservation affected the contractility of hiPSC-CMs relative to their fresh counterparts. Spontaneously contracting monolayers of hiPSC-CMs as well as single cells were recorded with a high-sampling rate camera (100 fps) and analysed using the automated open source software tool MUSCLEMOTION (Sala et al., 2018). We chose to examine cardiomyocytes from the hiPSC line LUMC20 due to this line showing electrophysiological differences between the two groups. Representative contraction traces of fresh and cryopreserved hiPSC-CM monolayers paced at 1 Hz are shown in **Fig. 4A**, while the contraction parameters analysed are illustrated in **Fig. 4B**. Because the recordings of frozen and non-frozen hiPSC-CMs could not be made concurrently, exposure conditions varied thereby precluding the comparison of contraction amplitudes between the two groups. Therefore, contraction traces are presented with a contraction amplitude normalized to the maximal value.

**Figure 4.**
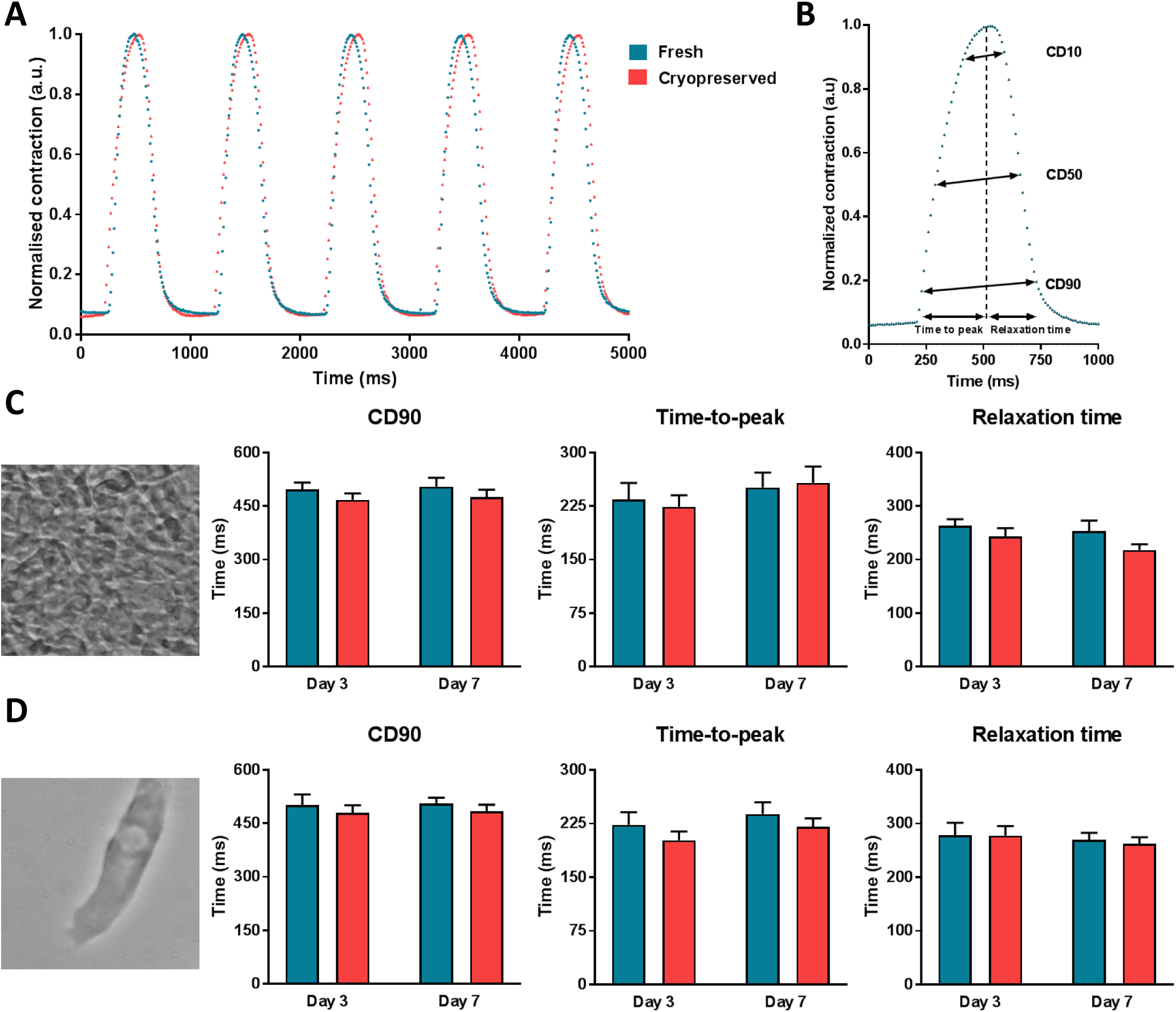
Contraction characteristics of non-frozen and cryopreserved hiPSC-CMs. **A)** Representative normalized contraction traces measured at 1 Hz for fresh and cryopreserved LUMC20 hiPSC-CMs seeded as a monolayer. **B)** Contraction parameters measured from a normalised contraction trace. **C & D)** Average data at 1 Hz for contraction duration at 90% peak amplitude (CD90), time-to-peak, and relaxation time for fresh and cryopreserved LUMC20 hiPSC-CMs measured in cells cultured either as monolayers (**C**) or single cells (**D**). Representative phase-contrast images are shown to the left of the bar graphs. For **C**, n=7 and 9 (day 3); n=8 and 9 (day 7) from 3 independent differentiations; For **D**, n=20 and 30 (day 3); n=28 and 28 (day 7) from 3 independent differentiations.

Contraction duration at 90% peak amplitude (CD90), time-to-peak, and relaxation time at day 3 and 7 post-replating were very similar between the fresh and cryopreserved cells in both hiPSC-CM monolayers and single cells (**Fig. 4C and 4D**). Congruently, no differences were observed for contraction duration (CD) at 10 and 50% peak amplitude (CD10 and CD50 respectively) values between the fresh and cryopreserved hiPSC-CMs (**Supplementary Fig. S4A**). Neither were differences detected when the fresh and cryopreserved hiPSC-CMs were stimulated at 2 Hz (**Supplementary Fig. S4B-4D**). Overall these results indicated that cryopreservation does not affect the contractility of the hiPSC-CMs.

## 3. Discussion

Advances in hPSC-CM generation and phenotyping have made them valuable as *in vitro* models for studying human heart development and cardiovascular disease, as well as for cardiac safety pharmacology, drug screening, and drug discovery (Brandão et al., 2017; Gerbin and Murry, 2015; Hartman et al., 2016; Magdy et al., 2018). While it is now possible to efficiently generate large quantities of relatively pure hPSC-CMs (Kempf and Zweigerdt, 2018), these procedures are time-consuming, laborious, and are still subject to variability in reproducibility between successive differentiations (Burridge et al., 2015; Kempf et al., 2014). Efforts to cryopreserve hPSC-CMs have led to improved consistency in functional assays with comparable results obtained when the same batch of cells are used even between different laboratories (Hwang et al., 2015; Millard et al., 2018). However, evaluation of the effect of cryopreservation on the *in vitro* characteristics of the hPSC-CMs are limited. For example, previous studies have either only assessed the cryopreserved hPSC-CMs (Hwang et al., 2015; Kim et al., 2011; Puppala et al., 2013), or comparisons were performed on cells immediately before and after thawing without assessing the recovery of the culture (Chen et al., 2015). Here we have directly compared the phenotype and physiological properties of frozen and non-frozen hPSC-CMs generated from the same differentiation experiment over a 10 day period post-replating. We have found that while cryopreservation can cause poorer replating of the hiPSC-CMs, this is not detrimental to their electrophysiological or contractile properties.

The (approximately two-fold) lower replating efficiency of the cryopreserved hPSC-CMs compared to non-frozen cells is possibly due to cryoprotectant solution (90% KSR, 10% DMSO) or the freezing protocol (−1°C/min to −80°C) used in this study, and further optimisation of these steps would likely improve the recovery rate. Several other reports have used foetal bovine serum (FBS) instead of KSR (Hwang et al., 2015; Kim et al., 2011). We chose not to include animal serum since it is undefined, can show batch variability and because all other maintenance and differentiation media used were serum-free. Other cryoprotectant solutions such as CryoStor CS-10 have also been used to cryopreserve hPSC-CMs (Chen et al., 2015; Xu et al., 2011); however we observed similar levels of recovery with this solution (data not shown). DMSO has been shown to be more cytotoxic to hiPSCs than other cryoprotective agents (Katkov et al., 2011), and so is also likely a cause of similar negative effects on hPSC-CMs. It is also apparent that the optimal cooling rate for cell survival varies depending on the cell type (Hunt, 2017). Therefore, further investigation of the most suitable cryoprotective agent and optimal cooling protocol for hPSC-CMs is warranted, in particular where these cells may be used therapeutically and large numbers of viable hPSC-CMs are required.

When the poorer replating efficiency of the cryopreserved hPSC-CMs was factored in to the replating density, there was no difference in the proportion of cardiomyocytes recovered when compared to non-frozen hPSC-CMs replated from the same differentiation experiment. Interestingly, in the hiPSC line LUMC20, we observed a trend towards a higher proportion of hPSC-CMs expressing the ventricular marker MLC2v following cryopreservation although this did not reach statistical significance. However, the cryopreserved cardiomyocytes from this cell line did display an increase in APD. These differences raise the possibility that cryopreservation could improve the functional maturation of hPSC-CMs. A similar observation was evident in hPSC-CMs stored under hypothermic conditions (+4°C), with the expression of several cardiomyocyte-specific genes including *MLC2V* significantly upregulated (Correia et al., 2016). The exact mechanism by which cold preservation could induce hPSC-CM maturation is unclear but hypothermic storage does increase caspase activity which has been demonstrated to promote the differentiation of mouse embryonic stem cell (ESC)-derived cardiac progenitors (Bulatovic et al., 2015). Alternatively, exposure of the hPSC-CMs to DMSO may also contribute to the enrichment in ventricular cardiomyocytes. Treatment of cultures with DMSO can downregulate *Oct-4* expression in mouse embryoid bodies as well as improve the differentiation of human ESC-derived pancreatic progenitors to terminal cell types (Adler et al., 2006; Chetty et al., 2013). In these studies, the concentration of DMSO was lower (<2% vs 10% in the cryoprotectant) but exposure was prolonged (1-2 days). Further studies are warranted to investigate whether cryopreservation or exposure of the hPSC-CMs to the cryoprotectant leads to enrichment of ventricular cardiomyocytes.

Importantly these results demonstrate that cryopreservation of hPSC-CMs does not adversely affect their functionality. We believe this is the first reported head-to-head comparative study that has systematically quantified APs and contraction kinetics in non-frozen and frozen hiPSC-CMs obtained from the same differentiation. Although the electrophysiological characteristics of fresh and cryopreserved cardiomyocytes derived from the human ESC line H7 have been analysed (Peng et al., 2010; Xu et al., 2011), these studies were performed using different experimental conditions, including the age of the hPSC-CMs measured, composition of electrophysiological solutions, recording conditions and classification of the cardiomyocytes. Therefore, it remains unclear whether the prolonged APD_90_ and slower V_max_ observed in the frozen hPSC-CMs in these studies was due to the cryopreservation or experimental setup.

In conclusion, we have shown that apart from replating efficiency, cryopreservation is not detrimental to hiPSC-CMs. Further modifications of the cryoprotectant solution composition as well as the freezing and thawing process will likely further improve this. The ability to freeze and recover functional hPSC-CMs, along with recent advances in efficient differentiation of hPSCs to cardiomyocytes, will likely contribute to improvements in standardisation. Not only will this enable the same batch of hPSC-CMs to be used in multiple assays and thereby allowing more direct comparisons for cardiac disease modelling, it will also overcome some of the challenges in using hPSC-CMs for large-scale screening of pharmacological compounds. Finally, facilitating distribution of identical batches hPSC-CMs between laboratories might lead to improved experimental reliability and robustness and contribute to addressing reproducibility issues in the field.

## 4. Material and Methods

### 4.1 hiPSC-CM differentiation

Subclones from two independent hiPSC lines (LUMC0020 and LUMC0099) were maintained in Essential 8 medium (Gibco). One day prior to differentiation (d-1), the hiPSCs were harvested using TrypLE Select (Gibco) and plated onto Matrigel-coated wells in Essential 8 medium containing Revitacell (1:200 dilution; Gibco) at 3.9 × 10^4^/cm^2^ in 12-well cell culture plates. The hiPSCs were differentiated into cardiomyocytes using the Pluricyte Cardiomyocyte Differentiation Kit (NCardia BV) according to the manufacturer’s instructions. The hiPSC-CMs were maintained in Medium C (NCardia) until day 20-21 of differentiation and then dissociated as previously described (Van Den Berg et al., 2014).

### 4.2 hiPSC-CM cryopreservation, storage and thawing

The hiPSC-CMs were cryopreserved in a freezing medium comprising 90% KSR (Gibco) and 10% DMSO. Cryovials containing ∼1 × 10^6^ cells in 300 µl freezing medium were rate-controlled (−1°C/minute) frozen to −80°C. Approximately 24 hours later, the vials were transferred and stored in liquid nitrogen (−196°C). For thawing, the vial was incubated at 37°C and the thawed cells transferred to a conical tube. Immediately thereafter, 1 ml of BPEL medium (Van Den Berg et al., 2014) was added dropwise (1 drop every 5 seconds), followed by ∼4.7 ml BPEL (1 drop every 2 seconds). Cell were precipitated at 250*g* for 3 min and resuspended in Medium C.

### 4.3 Replating of hiPSC-CMs

Fresh and thawed hiPSC-CMs were replated on Matrigel-coated glass coverslips, or in 24-well cell culture plates in Medium C supplemented with RevitaCell (1:100 dilution) at the densities indicated. Approximately 24 hours after plating the medium was refreshed, and subsequently every 2-3 days thereafter until the experiment was terminated.

### 4.4 Cell viability and replating efficiency

The viability of the dissociated hiPSC-CMs was determined by trypan blue staining (Davis et al., 2009). The viability and recovery of the replated cells was assessed at multiple time points using the CCK8 assay (Dojindo). For this, cells were seeded in multiple wells of a 24-well plate at varying densities (3.7-14.7 × 10^4^/cm^2^). For each timepoint, the cell culture medium was removed and replaced with 330 µl of reaction mixture (300 µl Medium C + 30 µl CCK8 reagent) per well. After 2.5h at 37°C, 100 µl from each well was transferred to a 96-well plate and the optical density (OD) at 450 nm measured using a Victor X3 microplate reader (Perkin Elmer). Reaction mixture added to a well without cells served as a blank control and was used to subtract the background fluorescence from the samples. The OD per 1 × 10^4^ cells was then calculated. After measuring, the CCK8 treated-cells were washed twice with cell culture medium and replaced with Medium C.

### 4.5 Flow cytometry

Cells were plated in 24-well cell culture plates at densities between 0.9 and 2.1 × 10^5^/cm^2^. Approximately 7 days post-seeding, the cells were dissociated using TrypLE Select and filtered to remove cell aggregates. The cells were incubated with a Viobility™ 405/520 fixable dye (Miltenyi Biotech) prior to fixation (FIX and PERM kit, Invitrogen) for subsequent exclusion of dead cells. Cells were co-labelled with cTnT (Vioblue-conjugated), MLC2a (APC- or FITC-conjugated), and MLC2v (PE-conjugated) antibodies (all Miltenyi). All antibodies were used at a concentration of 1:11 in permeabilization medium (medium B; Invitrogen). Data was acquired using a MacsQuant VYB flow cytometer (Miltenyi) and analysed with the software FlowJo (FlowJo, LLC).

### 4.6 Immunohistochemistry

The hiPSC-CMs were plated on glass coverslips at 0.7-1.3 × 10^4^/cm^2^ and fixed 6 days later using the Inside Stain Kit (Miltenyi) according to manufacturer’s instructions. Fixed cells were incubated with α-actinin (1:250; Sigma) and myosin heavy chain (1:50; Miltenyi) antibodies. The primary antibodies were detected with AF594- (1:200; Life Technologies) and Vio515-(1:100; Miltenyi) conjugated secondary antibodies, respectively. All antibodies were diluted in permeabilization medium and incubated for 10 minutes. Cells were stained with 4’,6 Diamidino-2-Phenylindole (DAPI) (0.3 µM) for 5 minutes. Images were captured using a confocal laser scanning microscope SP8 (Leica).

### 4.7 Action potential measurements

The hiPSC-CMs were plated on 10 mm glass coverslips at 1.0 × 10^4^/cm^2^ and APs were recorded 8-10 days later using the perforated patch-clamp technique and an Axopatch 200B amplifier (Molecular Devices). Single cells with spontaneous contractions (indicating the viable state of the cells) were selected. Using dynamic clamp, an in silico 2 pA/pF I_K1_ with Kir2.1 characteristics was injected to obtain quiescent cells with a close-to-physiological RMP as previously described (Meijer van Putten et al., 2015). Data acquisition, voltage control, and analysis was performed using custom made software and the potentials were corrected for the calculated liquid junction potential (Barry and Lynch, 1991). The patch pipettes, estimations of cell membrane capacitance, and filtering and digitizing settings were as described previously (Verkerk et al., 2017).

Cells were superfused with modified Tyrode’s solution (36 ± 0.2°C) containing (in mM): 140 NaCl; 5.4 KCl; 1.8 CaCl_2_; 1.0 MgCl_2_, 5.5 glucose; and 5.0 HEPES. (pH was set at 7.4 using NaOH). Pipettes were filled with a solution containing (in mM): 125 K-gluconate; 20 KCl; 5.0 NaCl; 0.44 amphotericin-B; and 10 HEPES (pH 7.2; KOH). APs were elicited at 1 Hz by 3 ms, ∼1.2x threshold current pulses through the patch pipette, and analysed for RMP, V_max_, APA, APD_20_, APD_50_, and APD_90_. Parameters from 13 consecutive APs were averaged.

### 4.8 Contraction measurements

The contraction of single cells and monolayers of hiPSC-CMs was determined by seeding cells on 10 mm glass coverslips at a density of 1.0 or 12.7 × 10^4^/cm^2^ respectively. At day 3 and 7 post plating, coverslips were transferred into a bath superfused with modified Tyrode solution at 37°C. The single cells and monolayers were paced at 1 and 2 Hz using an external field stimulator. A 10s movie was recorded using a DCC3240M camera (Thorlabs) with a sampling rate of 100 frames per second (fps). From the normalized contraction traces, the CD10, CD50, CD90, time-to-peak, and relaxation time were calculated using the automated, open source software tool MUSCLEMOTION21.

### 4.9 Statistical data analysis

All data are presented as mean ± SEM. Statistical tests performed are listed in the Results section or in the Fig. legends. Differences were considered statistically significant at p<0.05. Analyses were conducted with Graphpad Prism 7 software.

## Acknowledgments

We thank Leon Tertoolen and Berend van Meer for assistance with the MUSCLEMOTION software and Jelle Goeman for statistical advice. This work was supported by a Starting Grant (STEMCARDIORISK) from the European Research Council (ERC) under the European Union’s Horizon 2020 Research and Innovation programme [H2020 European Research Council; grant agreement #638030], and a VIDI fellowship from the Netherlands Organisation for Scientific Research [Nederlandse Organisatie voor Wetenschappelijk Onderzoek; ILLUMINATE; #91715303].

## Author contributions

L.vd.B. designed and carried out the majority of the experiments, performed the analysis and wrote the manuscript. K.B., C.G. and M.M. performed the hiPSC-CM differentiations and assisted in cryopreserving the hiPSC-CMs. C.L.M. and A.O.V. provided scientific support and edited the manuscript. R.P.D. conceived and supervised the project, designed and carried out experiments and wrote the manuscript.

