## Supplementary Data for "Cryopreservation of human pluripotent stem cell-derived cardiomyocytes is not detrimental to their molecular and functional properties"

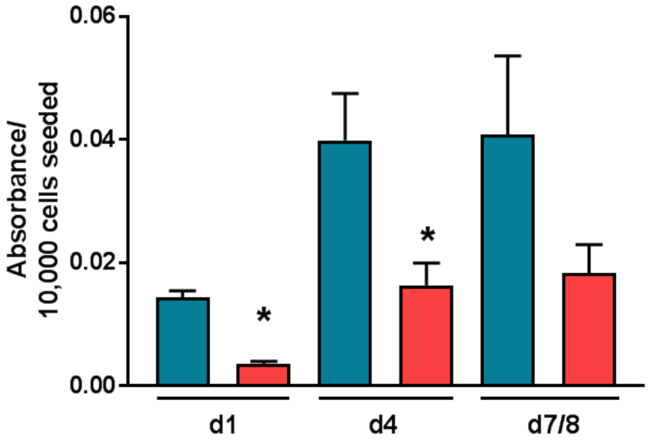


**Figure S1** Recovery of non-frozen (blue) and cryopreserved (red) LUMC99 hiPSC-CMs at day 1, 4 and 7/8 post-replating. n=4-6 from 2 independent differentiations. * indicates statistical significance (day 1 p<0.001, day 4 p=0.014, unpaired *t*-test)

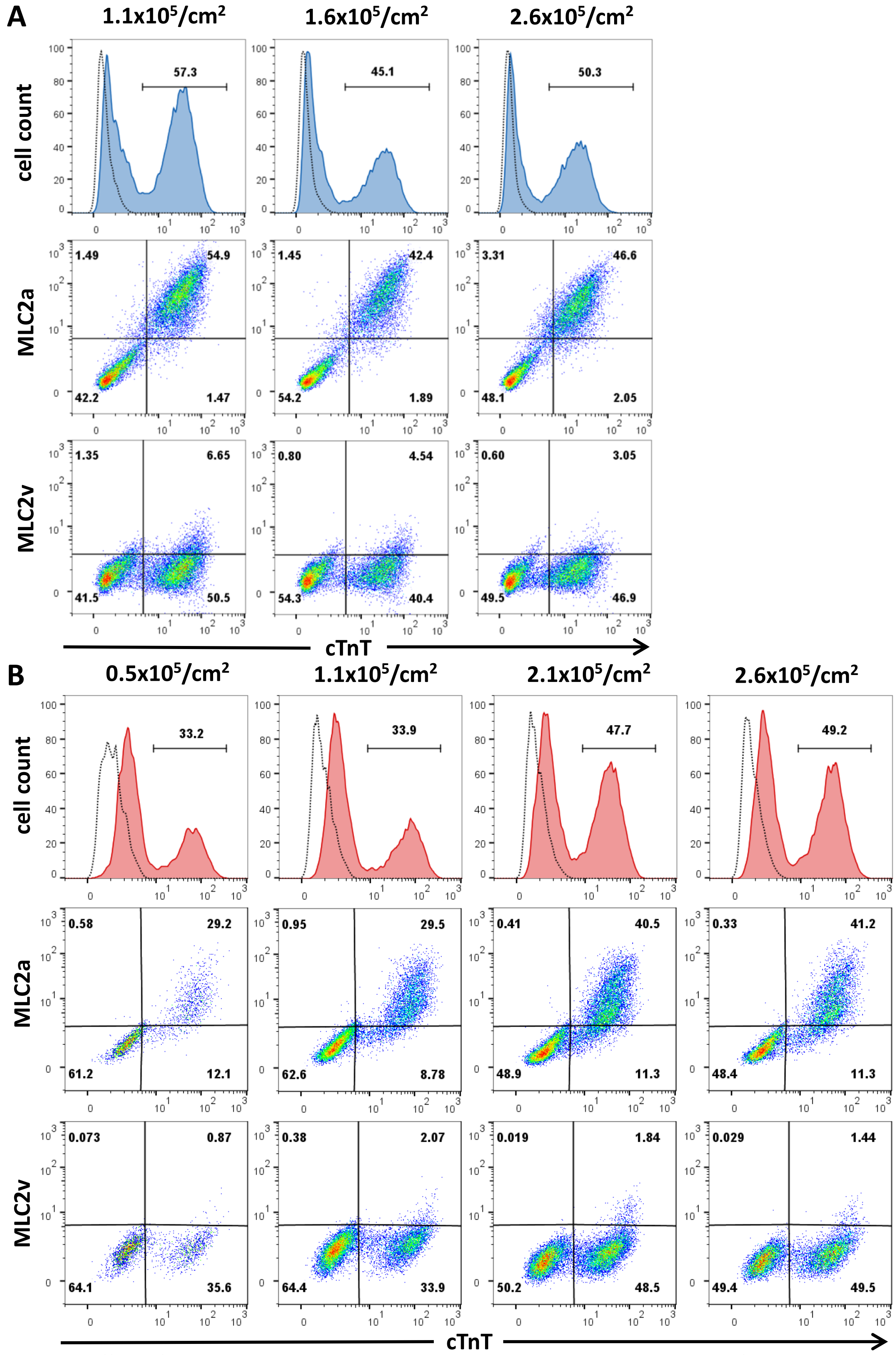


**Figure S2** Effect of replating density on cTnT, MLC2a and MLC2v expression in non-frozen and cryopreserved hiPSC-CMs. Flow cytometry analysis indicated replating density had little effect on the expression of these markers when the cardiomyocytes had not been frozen (**A**), however for cryopreserved hiPSC-CMs (**B**), higher replating densities resulted in an increased proportion of cells expressing cTnT and MLC2a. Values above plots indicate the density the hiPSC-CMs were replated. Top row depicts histogram plots of cTnT expression, while remaining rows depict bivariate density plots of MLC2a/cTnT (*middle row*) and MLC2v/cTnT (*bottom row*). Numbers inside the plots are the percentage of cells within the gated region. Dotted lines represent a control cTnT$^-$ population.

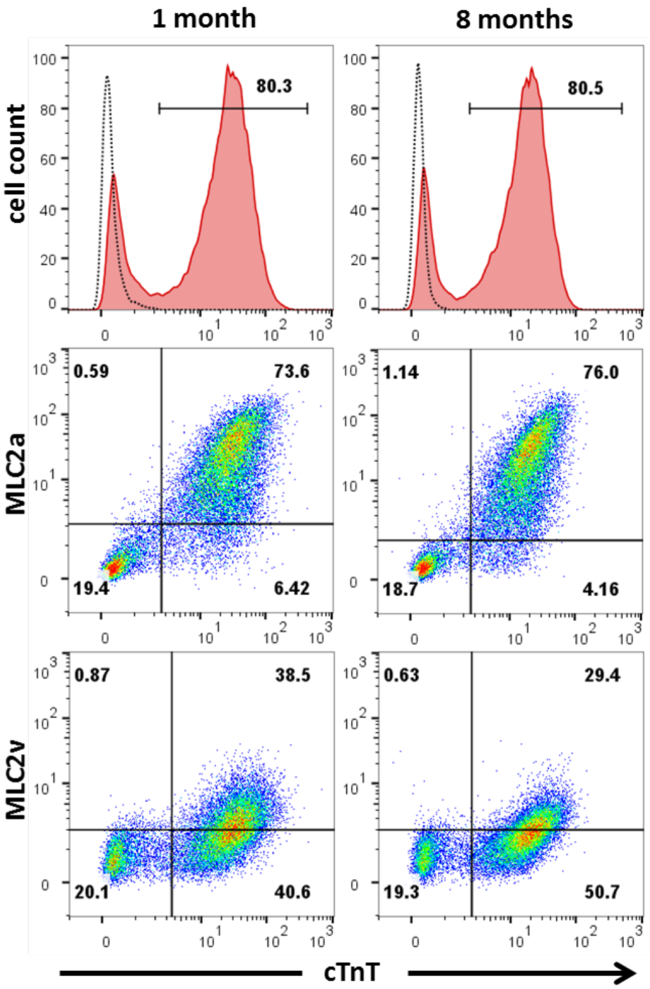


**Figure S3** No differences in cTnT, MLC2a and MLC2v expression were detected by flow cytometry analysis in LUMC20 hiPSC-CMs that had been cryopreserved for 1 month (*left column*) or for 8 months (*right column*) when replated at similar densities. Top row depicts histogram plots of cTnT expression, while remaining rows depict bivariate density plots of MLC2a/cTnT (*middle row*) and MLC2v/cTnT (*bottom row*). Numbers inside the plots are the percentage of cells within the gated region. Dotted lines represent a control cTnT$^{-}$ population.

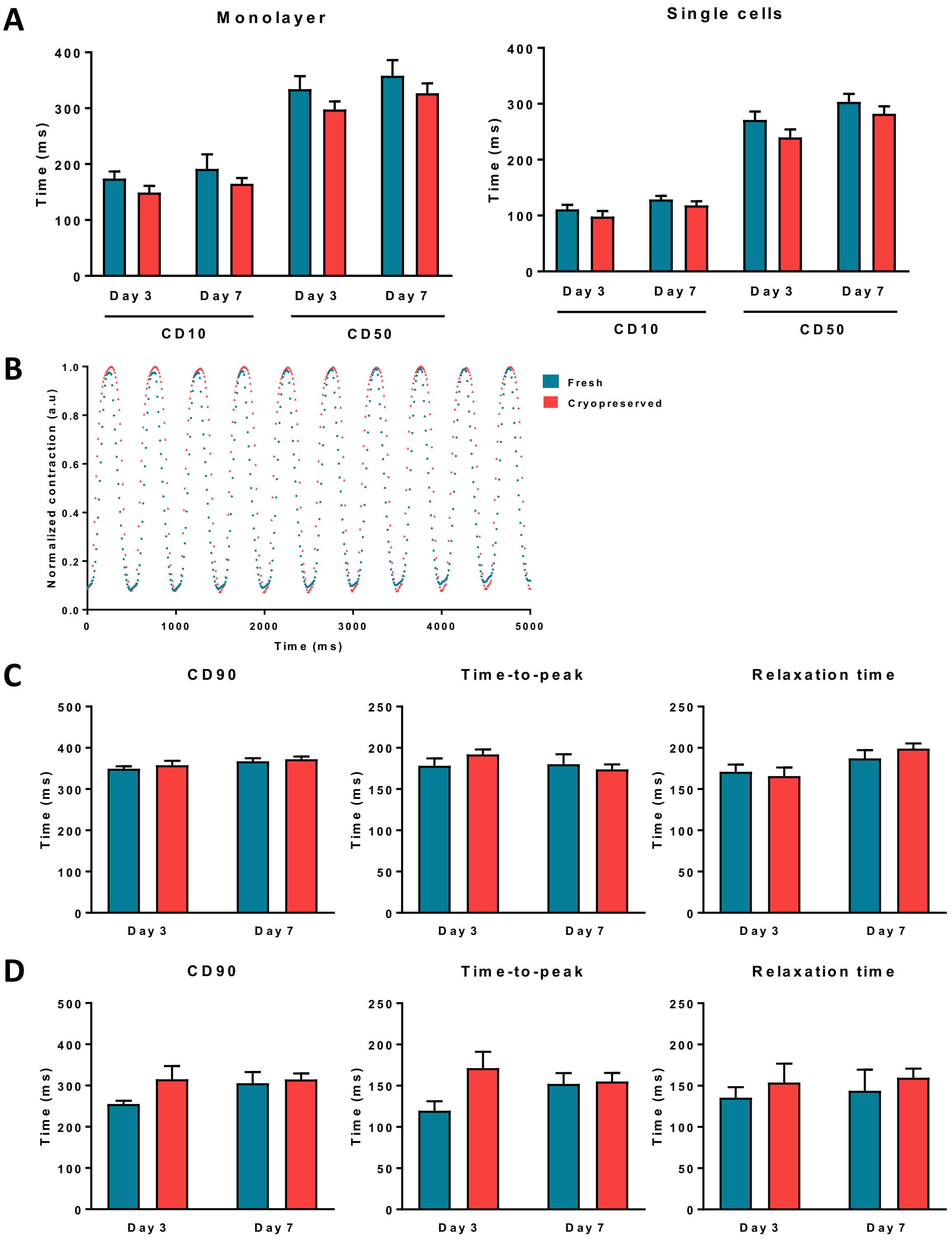


**Figure S4** **A)** Average data at 1 Hz for contraction duration at 10% and 50% from peak (CD10, CD50) for fresh (blue) and cryopreserved (red) LUMC20 hiPSC-CMs seeded as a monolayer (left) or as single cells (right). **B)** Representative normalized contraction traces measured at 2 Hz for fresh and cryopreserved LUMC20 hiPSC-CMs seeded as a monolayer. **C & D)** Average data at 2 Hz for CD90, time-to-peak and relaxation time for fresh and cryopreserved LUMC20 hiPSC-CMs measured in cells cultured either as monolayers (**C**) or single cells (**D**). For **C**, n=6 and 9 (day 3); n=7 and 9 (day 7) from 3 independent differentiations; For **D**, n=4 and 7 (day 3); n=5 and 10 (day 7) from 2 independent differentiations.
